# The Spectral Sensitivity of the Neurons Mediating Black and White

**DOI:** 10.1101/829051

**Authors:** Sara S. Patterson, Maureen Neitz, Jay Neitz

## Abstract

Our percepts of black and white are not equally strong for all monochromatic lights across the spectrum, but instead have a spectral tuning defined by they ways in which their neural substrates process the outputs of three univariant cone photoreceptors. The neurons mediating black and white and how they combine the cone outputs remain controversial but growing evidence indicates cone-opponent midget ganglion cells are involved. The paradoxical implications of having “chromatic” neurons mediate what is traditionally assumed to be a role of “achromatic” neurons remain unresolved. Here, we investigate whether midget ganglion cells can account for the variation in perceived saturation with wavelength.

## 1. INTRODUCTION

Our experience of color can be defined by three parameters: hue, brightness and saturation. A fundamental question in color vision research is how these perceptual features relate to the physical attributes of incoming light and how they are represented by neurons within the visual system. Of these three parameters, saturation has been the most difficult to link to an underlying neural substrate.

Saturation is the perceived distance of a color from white, gray or black (black-white) [1]. For example, a patch of monochromatic blue light against a relatively dark background appears very saturated, producing little or no white sensation. Between unique blue and equal energy white, the perceived color becomes increasingly desaturated as the strength of the white sensation increases relative to the blue hue sensation. The same applies to the perceived distance from black for colored patches against lighter backgrounds – navy appears desaturated because the black sensation is strong relative to the blue hue sensation. When matched for brightness, colors have two components - hue and black-white - with the relative strengths of these two components defining the perceived saturation [2, 3].

The relative amounts of hue and black-white perceived in monochromatic lights vary reliably as a function of wavelength. Trichromats universally agree that of the four fundamental hues (red, green, blue and yellow), yellow is the most desaturated (i.e. contains the most white) and blue is the most saturated. Red is perceived as slightly less saturated than blue while green is perceived as slightly more saturated than yellow. This phenomenon is quantified in classic saturation discrimination experiments, such as Figure 1A and Figure 1D. The resulting saturation discrimination function is interpreted as reflecting the relative activity of the neural substrates for hue and black-white as a function of wavelength. However, the identities of the neural substrates for both hue and black-white remain highly controversial.

**Fig. 1.**
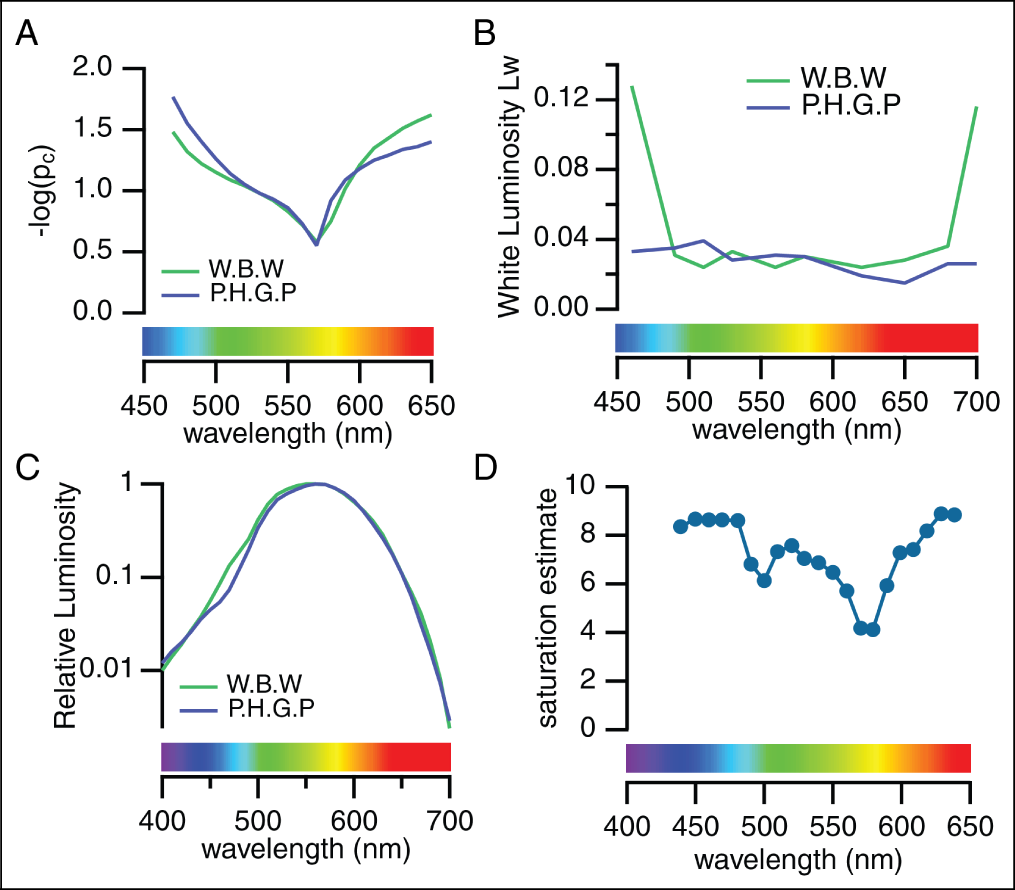
**(A)** Saturation discrimination curves measured as the “first step from white” from Wright & Pitt (1937) [12]. **(B)** Saturation discrimination curves measured as the “first step from the spectrum” from Wright & Pitt (1935) [13]. **(C)** Spectral luminosity functions, *V*(*λ*), from the same two observers, obtained from Wright & Pitt (1935) [13]. **(D)** Saturation scaling experiment from Jacobs (1967) [14].

Classic models fitting the saturation discrimination function assumed black-white was mediated by neurons summing L- and M-cones (*L* + *M*) [4–7]. However, there is now growing evidence that our percepts of black and white are instead mediated by spectrally-opponent (*L−M*) midget ganglion cells [8–11]. The purpose of this study is to determine whether the spectral sensitivities of midget ganglion cells can account for the saturation discrimination function.

## 2. METHODS

### A. Saturation discrimination experiments

There are several different experimental paradigms for measuring saturation as a function of wavelength and they produce different results (Figure 1A, 1B, 1D). Though undoubtedly related to saturation, the results of some paradigms do not perfectly match our perceptual experience of saturation [15]. Though the paradigms are sometimes grouped together for analysis and interpretation [4], here we will examine each paradigm individually.

#### A.1. First Step from White Paradigm

Our primary focus will be modeling a classic experimental paradigm for quantifying the wavelength dependence of saturation that measures the “first step from white”[16] or the “chromatic threshold with respect to white”[15] (Figure 1A). A broadband light (perceived as white) is presented to the entire field while a monochromatic spectral light is added to just one half of the field (Figure 4A). The strength of the spectral light is increased until just noticeably different from the broadband light. In other words, until a just noticeable difference in saturation occurs. Importantly, the two lights are also matched for brightness. Then the least perceptible colorimetric purity (*p*_*c*_) can be obtained by:

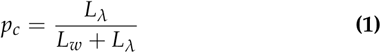

where *L*_*λ*_ is the intensity of the spectral light and *L*_*w*_ is the intensity of the broadband light. This metric expresses saturation independent of both the intensity of the broadband light and the wavelength of the spectral light [15]. By convention, the results are plotted as the inverse log, −*log*(*p*_*c*_), so that the y-axis reads out as a measure of saturation as a function of wavelength. In other words, Figure 1A can be read as saying the participant had to increase the intensity of the 570 nm light far more than the intensity of the 470 nm light to produce a discriminable change in saturation, defined as a just-noticeable-difference in hue.

#### A.2. First Step from the Spectrum Paradigm

The “first step from the spectrum” experiments in Figure 1B measure the minimal amount of a broadband light that must be added to a monochromatic spectral light to produce a just-noticeable decrease in saturation. While initial studies found relatively flat sensitivity across the visual spectrum [12, 15], subsequent work reported more wavelength-dependence [17] and an interesting relationship with mean light level not seen in the “first step from white” paradigm [16]. Additional studies will be necessary to address these inconsistencies so we do not attempt to fit these results here.

An important point is that both the “first step from white” and the “first step from the spectrum” rely on threshold changes in saturation, yet the results are strikingly different (compare Figure 1A to Figure 1B). However, the way each experiment defines the saturation threshold is also different. The “first step from white” measures a threshold hue percept while the “first step from the spectrum” measures a threshold white percept.

#### A.3. Saturation Scaling Paradigm

The saturation scaling paradigm in Figure 1D asks participants to rate the saturation of equiluminant spectral lights on a fixed scale [14, 18, 19]. Saturation scaling is distinct from the “first step from white” and “first step from the spectrum” paradigms because it is not a threshold measurement but rather a direct judgment of perceived saturation.

### B. Model

#### B.1. Cone-opponent midget ganglion cells

Traditionally, accounts of the physiological basis of saturation have assumed contributions from spectrally-opponent neurons for hue and spectrally non-opponent neurons for black-white [23] (but see De Valois & Marrocco, 1973 [24]). Here we took a different approach and limited our model to the midget ganglion cell subtypes proposed to be involved in black and white (Figure 2). Each L- and M-cone photoreceptor in the central retina is represented twice in the retinal output, by both an ON and OFF midget ganglion cell [25]. In addition, a controversy regarding S-cone midget ganglion cells [26, 27] was recently resolved and it was confirmed that each S-cone has an OFF midget ganglion cell, but not an ON midget ganglion cell [28]. Midget ganglion cells have center-surround receptive fields that provide both spatial and spectral opponency (Figure 2, reviewed in Patterson et al., 2019 [9]).

**Fig. 2.**
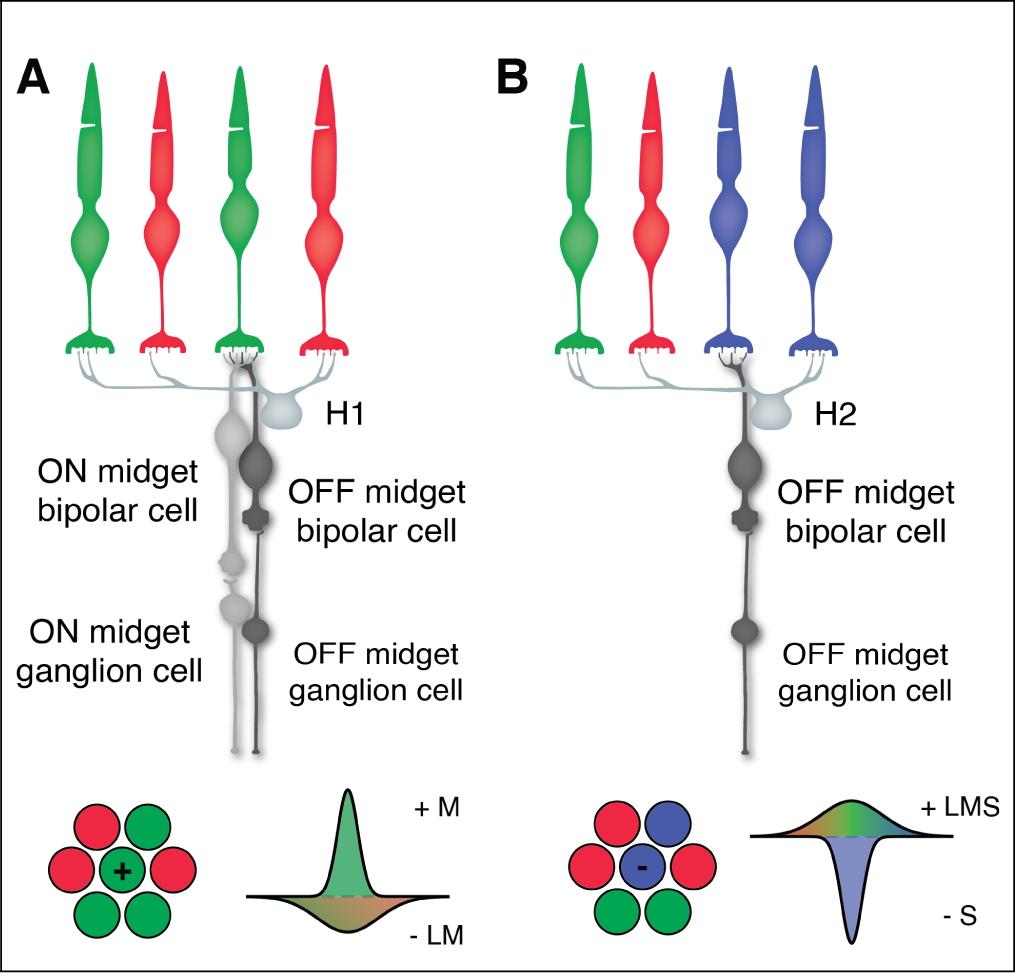
Spatial and spectral receptive fields of midget ganglion cell subtypes. **(A)** Circuitry for M-ON and M-OFF midget ganglion cells. The circuitry for L-ON and L-OFF midget ganglion cells is identical, but with an L-cone directly contacting the midget bipolar cells and forming the center receptive field (not pictured). **(B)** Circuitry for S-OFF midget ganglion cells. There is no S-ON midget ganglion cell. The photon catch in the center cone is compared to the photon catch in surrounding cones provided by H1 or H2 horizontal cell feedback [20–22]. Bottom: Illustration of the cone inputs and sensitivity profiles for M-ON and S-OFF midget ganglion cell receptive fields. The polarity of the center cone is marked with ‘+’ or ‘-’ for ON and OFF midget ganglion cells, respectively.

The outputs of midget ganglion cell center-surround receptive fields are a measure of the relative photon catch in the center and surround receptive fields. Their outputs vary as a function of wavelength because the cone inputs to the center and surround receptive fields differ and each cone’s response is univariant, confounding changes in wavelength with changes in intensity [29, 30]. Here, we modeled the wavelength-dependence of the midget ganglion cell’s output as the difference between the spectral sensitivities of the center and surround receptive fields (Figure 3) [31, 32].

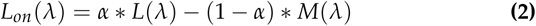

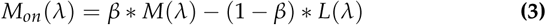

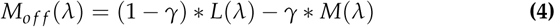

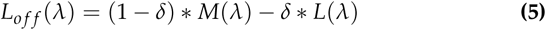

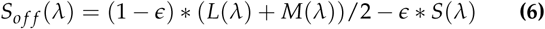

where *L*(*λ*), *M*(*λ*) and *S*(*λ*) represent the L-, M- and S-cone spectral sensitivities, respectively, corrected for optical density and macular pigment. The Neitz photopigment template was used, assuming peak sensitivities of 559, 530 and 419 nm and optical densities of 0.3, 0.3 and 0.35 for the L-, M- and S-cones, respectively [33]. The center-surround balance is represented by *α*, *β*, *γ*, *δ* and *e* for M-ON, L-ON, M-OFF, L-OFF and S-OFF midget ganglion cells, respectively. The output of each model midget ganglion cell was rectified to simulate the nonlinearity imposed by the spike threshold of a ganglion cell. The equations for the five spectral subtypes of midget ganglion cells are schematized in Figure 3.

**Fig. 3.**
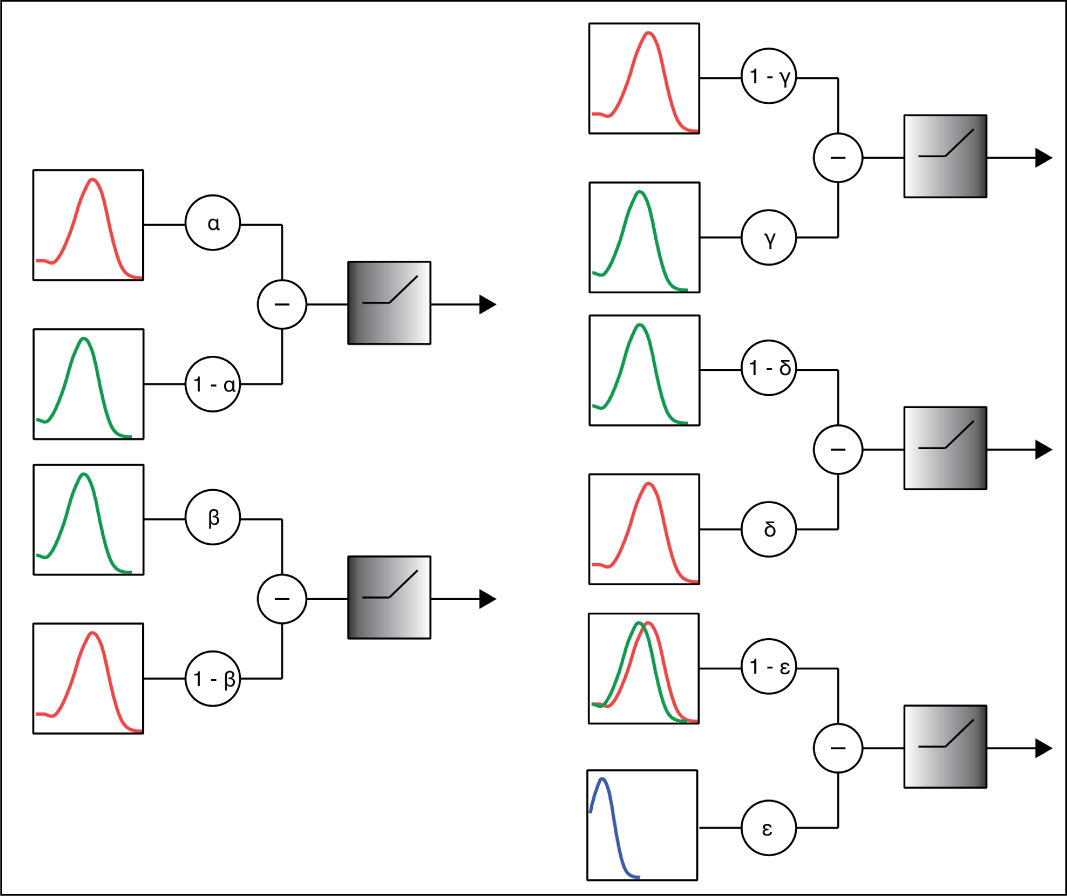
The output of a midget ganglion cell is a measure of relative photon catch in the center and surround receptive fields. However, this varies as a function of wavelength because the center and surround receptive fields have different spectral sensitivities. To capture this, we modeled the outputs of the five spectral subtypes of midget ganglion cell as a function of wavelength. L-ON and M-ON midget ganglion cells are on the right. L-OFF, M-OFF and S-OFF midget ganglion cells are on the left. Cone spectral sensitivities for the center and surround receptive fields are weighted with opponent gain parameters, subtracted from each other then passed through a rectifying nonlinearity. The final saturation discrimination models combine these five independent retinal output pathways in various ways to account for the “first step from white” saturation discrimination function.

While these equations appear to maximize spectral opponency, the presence of, for example, L-cones in the surround of the L-OFF midget ganglion cell, can be accounted for by decreasing the *α* parameter. This parameter can be varied to account for the spatial structure of the stimulus (discussed in the **Results**). This term also reduces the number of assumptions we need to make about crucial unknown variables, such as the relative strength of the center and surround receptive fields and the participant’s L:M-cone ratio.

#### B.2. Saturation models

Classic saturation models proposed color saturation was mediated by the ratio of activity in the neural substrates for hue and black-white [4–6, 23]. The spectral sensitivities of the neural substrates for hue were assumed to match the red-green and blue-yellow opponent response functions obtained with hue cancellation [34, 35], while the neural substrates for black-white were assumed to match the spectral luminosity function, *V*(*λ*), obtained with heterochromatic flicker [36, 37]. Our approach is distinct as we instead constrained our model to the features of midget ganglion cell receptive fields established through anatomical and physiological experiments [20–22, 25, 26, 28]. The resulting midget ganglion cell spectral sensitivities are distinct from the opponent response functions and the spectral luminosity function obtained through psychophysics experiments. Because the output of each midget ganglion cell subtype is rectified, the models developed in the **Results** (Eq. (7) and Eq. (8)) are linear combinations of the spectral sensitivities of the outputs of midget ganglion cells, not the L-, M- and S-cone spectral sensitivities, as in other color vision models that account for the saturation discrimination function [4–6].

We adopted three standard assumptions common to models fitting the saturation discrimination function. First, that there is a set of neurons whose activity directly determines the strength of our fundamental percepts of red, green, blue, yellow, black and white. We refer to these as the neural substrates of hue and black-white. A second assumption is that ON cells mediate percepts of white while OFF cells mediate percepts of black [8, 38, 39]. Third, we assume that the characteristics of the underlying neural substrates of hue and black-white (in our case, the midget ganglion cells) can be combined linearly to account for the saturation discrimination function. Linear combinations are generally considered valid for experiments measuring threshold discriminations [5], such as the “first step from white” paradigm.

#### B.3. Model Fitting Procedure

We obtained saturation discrimination data for the “first step from white” paradigm from published tables [2, 12, 15, 40]. The scaling data was extracted from a published figure [14]. The models were fit to each individual saturation discrimination curve dataset in MATLAB (Mathworks) using the lsqcurvefit function. The y-axis scaling varied between experiments, so a scaling parameter, *θ*, was included. The resulting curves were translated along the y-axis to align with the minimum of each saturation discrimination dataset. Additional details about each model are discussed as they are introduced in the **Results**.

## 3. RESULTS

### A. Development of a Midget Ganglion Cell Model for the Saturation Discrimination Function

We began by asking whether midget ganglion cells alone could account for the results of the “first step from white” paradigm, as this is the classic method for obtaining saturation discrimination functions.

#### A.1. Model 1: LM-ON midget ganglion cells

Because the saturation discrimination experiments directly involve the perceptual experience of white, we began by asking whether ON midget ganglion cells could account for the saturation discrimination function. While it was clear their spectral sensitivities could not provide a full account, we asked whether they could at least explain a particularly salient feature: the sharp minimum around 570 nm.

We modeled the spectral sensitivity of L-ON and M-ON midget ganglion cells using Eq. (2) and Eq. (3), respectively. When the center and surround receptive fields were equally weighted (*α* = *β* = 0.5), the peak in white percepts at 570 nm was not predictable from the spectral sensitivities of the ON midget ganglion cells (Figure 4C).

**Fig. 4.**
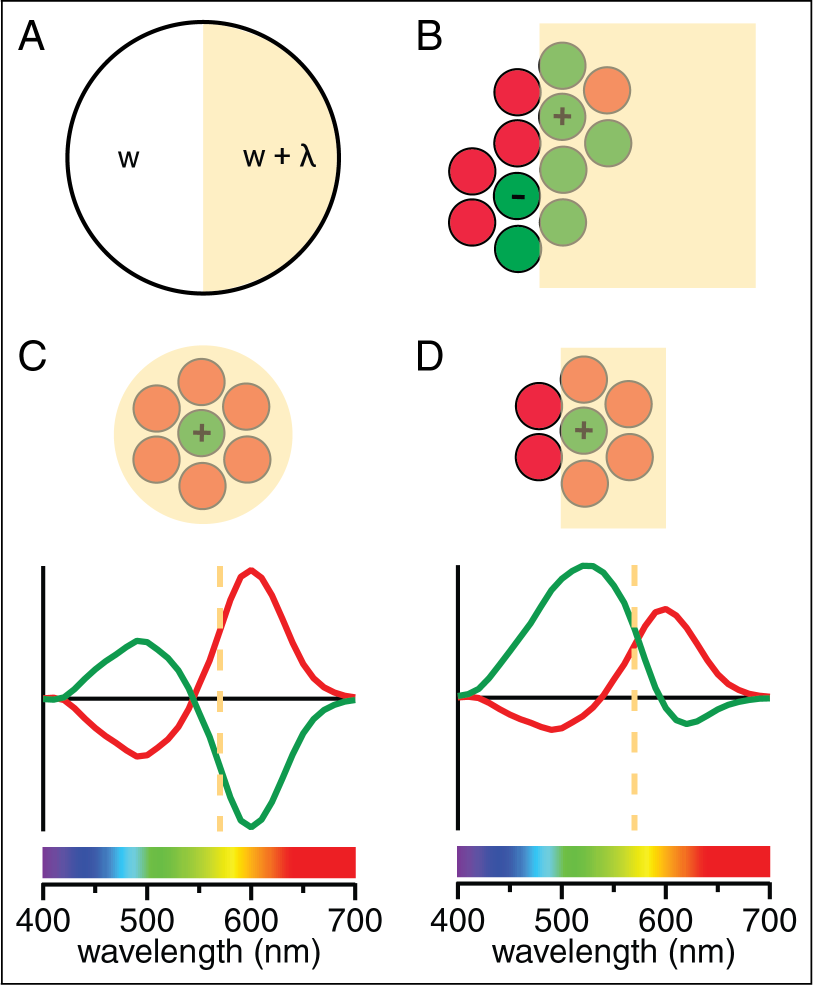
**(A)** Example stimulus for a saturation discrimination stimulus experiment. **(B)** Cone inputs to L/M midget ganglion cells at the edge of the test light. An ON-center cell responds best at the inner edge of the test light while an OFF-center cell responds best outside the test stimulus. Note that the model also includes a midget ganglion cell of the opposite polarity at each center cone. **(C, D)** Top: Illustration of full-field and edge stimuli superimposed on the receptive field of a sample M-ON midget ganglion cell. Bottom: Corresponding L-ON and M-ON midget ganglion cell spectral sensitivities before rectification. Yellow dashed line at 570 nm.

The approach in Figure 4C considers only the spectral tuning of the cone inputs to midget ganglion cells and does not account for the spatial arrangement of those cone inputs. The combination of spectral and spatial opponency in midget ganglion cells means wavelength and spatial structure are confounded. Figure 4C overlooks the midget ganglion cell’s spatial opponency and essentially isolates the spectral opponency by modeling the spectral response of a neuron to a spatially-homogeneous monochromatic stimulus. Yet there is considerable evidence that our perception of hue and black-white is instead defined by the changes in intensity and wavelength that occur at borders [41, 42]. The saturation of the spectral light is likely defined by the receptive fields of neurons located along the edge of the stimulus and the saturation of the interior is set by “filling in” at the level of the cortex. Consistent with this idea, the center-surround receptive fields of midget ganglion cells are ideal edge detectors and encode spatial contrast, such as that occurring at a border [9, 43]. As shown in Figure 4D, if the receptive field is located at an edge and only some of the surround is stimulated (or the weight parameters *α* and *β* are adjusted), the contributions of both types now could sum to signal white at 570 nm. Because spatial and spectral information are confounded in the midget ganglion cell’s output, changes in the spatial structure of the stimulus will alter the midget ganglion cell’s spectral sensitivity.

#### A.2. Model 2: LM-ON and LM-OFF midget ganglion cells

The L-ON and M-ON midget ganglion cell spectral sensitivities in Figure 4D can contribute to peak white sensations at 570 nm but cannot account for the steep increases in saturation surrounding 570 nm. In other words, the strength of white percepts in the middle wavelengths is too broad and would need to be attenuated to fully account for the saturation discrimination function. We next investigated whether including OFF midget ganglion cells could provide a full account by asking whether the saturation discrimination function could be fit by a model combining the outputs of LM-ON and LM-OFF midget ganglion cells (but without S-OFF midget ganglion cell contribution).

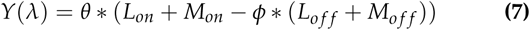

where *φ* represents the gain of the OFF midget ganglion cells and *θ* is the scaling parameter introduced in the **Methods**. Note that opponency between the ON and OFF cells is not strictly enforced by the model as in Eq. (2)–Eq. (6). The OFF gain parameter, *φ*, only determines the strength of the OFF cells relative to the ON cells, which were set to an arbitrary gain of 1.

The results (red curve in Figure 5A) demonstrate excellent fits to the middle and long wavelength regions of the saturation discrimination function. L vs. M cone-opponency in midget ganglion cells provides a clear explanation for the sharp minimum at 570nm. However, the model fails to account for the slope of increasing saturation for short-wavelength lights and instead predicts monochromatic blues and reds would be equally saturated.

**Fig. 5.**
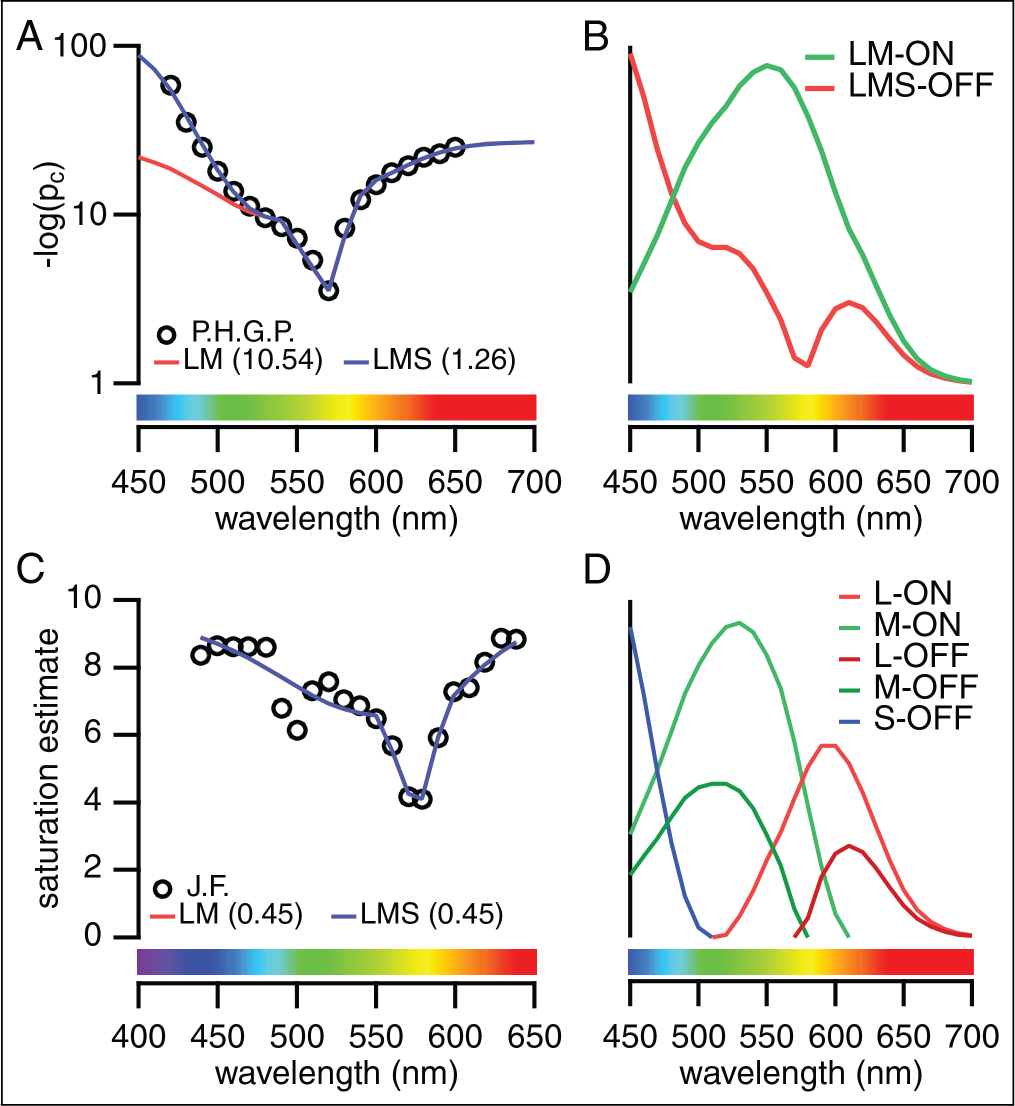
**(A)** Midget ganglion cell models with (blue curve) and without (red curve) the S OFF midget ganglion cell fit to the saturation discrimination data for one observer, P.H.G.P., from Wright & Pitt (1937) [12]. The root mean square error is shown in parentheses. **(B)** Models fit to saturation scaling data for one observed, J.F., from Jacobs (1967) [14]. **(C)** Summed spectral sensitivities for ON and OFF midget ganglion cell subtypes for the fit in **A**. **(D)** Spectral sensitivities for each spectral subtype of midget ganglion cell for the fit in **A**.

#### A.3. Model 3: LM-ON and LMS-OFF midget ganglion cells

Accounting for the short-wavelength component of the saturation discrimination function appears to require contribution from S-cones. There has been considerable controversy regarding the existence of S-OFF midget ganglion cells and these neurons have not been incorporated into models of visual perception. We recently provided both anatomical and physiological confirmation of the existence of S-OFF midget ganglion cells in the primate central retina [28].. Briefly, we confirmed a single OFF midget bipolar cell contacted each S-cone, which in turn, contacted a single OFF midget ganglion cell. We also confirmed that, like the more common L vs. M-cone midget ganglion cells, S-OFF midgets have a center-surround receptive field consistent with a role in spatial vision.

To determine whether the S-OFF midget ganglion cells are necessary to account for the saturation discrimination function, we added an S-OFF term (Eq. (6)) to the previous model.

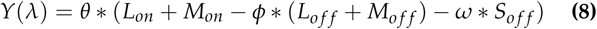

The S-OFF term was given a separate gain control (*ω*) from the L/M-OFF terms to allow the model to account for the sparsity of S-cones and the potential for their signals to be amplified downstream [44, 45].

Including the S-OFF midget ganglion cell allowed our model to fit the short-wavelength region of the saturation discrimination function (blue curve in Figure 5A). To provide intuition on how the model worked, the spectral sensitivities of the outputs of each midget ganglion cell subtype are shown in Figure 5D and the ON and OFF subtypes summed together are shown in Figure 5B. Our model provided excellent fits to many “first step from white” datasets (Figure 6) [2, 12, 15, 46].

**Fig. 6.**
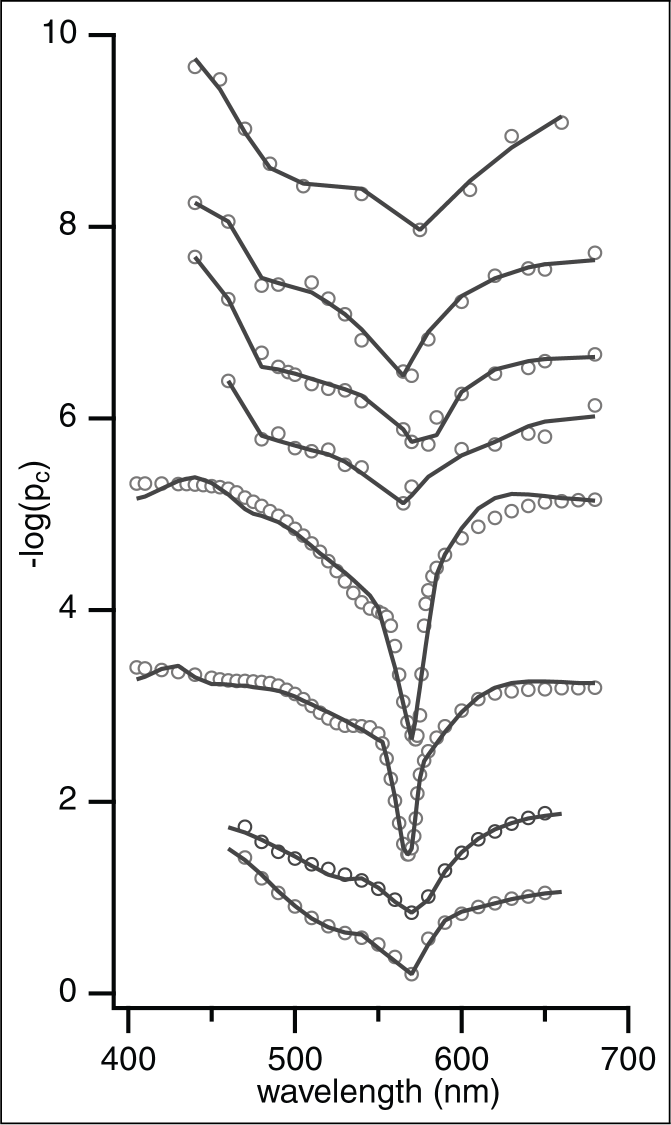
Model fits (grey) to classic saturation discrimination datasets (black circles). From top to bottom: D.P. from Purdy (1931) [15], W.J.M., L.C.M. and F.L.W., from Martin et al (1933) [2], I.G.P. and F.G.B. from Priest & Brickwedde (1938) [46], W.D.W. and P.H.G.P. from Wright & Pitt (1937) [12].

### B. Application of Model to the Saturation Scaling Paradigm

We next asked whether our models could account for the results of the saturation scaling paradigm. Some past models have grouped scaling and “first step from white” saturation discrimination functions together as being roughly similar [4]. However, there are several reliable differences between results obtained by the two paradigms (compare Figure 1A and Figure 1D). The clearest difference is the strong decrease in saturation at 490-500 nm obtained with the scaling paradigm and the steeper short-wavelength slope obtained with the “first step from white” paradigm.

Attempting to fit the results of the saturation scaling paradigm with our models was not as successful but revealed two interesting features (Figure 5C). First, our model cannot account for the dip in saturation at 490-500 nm or the shallow slope in the short wavelength region. Second, the S-OFF midget ganglion cell parameter did not alter the fit, indicating this parameter was necessary for the “first step from white” paradigm, but not for the scaling paradigm.

### C. Saturation Discrimination in Dichromats

Our model demonstrated that the L vs. M cone opponency in midget ganglion cells can explain the sharp minimum in saturation around 570 nm. Additional support for this conclusion can be seen in the saturation discrimination functions of dichromats (Figure 7A) [40, 47–49]. The effect of a missing L-, M- or S-cone on the spectral luminosity function, *V*(*λ*) is predictable - the peak shifts towards the remaining long-wavelength cone types (Figure 7B and Figure 7D). However, the effect of dichromacy is more complex for saturation discrimination functions. The sharp decrease in saturation around 570 nm is absent for protanopes and deuteranopes (Figure 7A), but remains desaturated for tritanopes (Figure 7C) [48]. Interestingly, for deuteranopes and protanopes, the saturation discrimination functions are closer to matching our intuition of the ideal white (i.e. equal sensitivity at all wavelengths) than the spectral luminosity functions (Figure 7A).

**Fig. 7.**
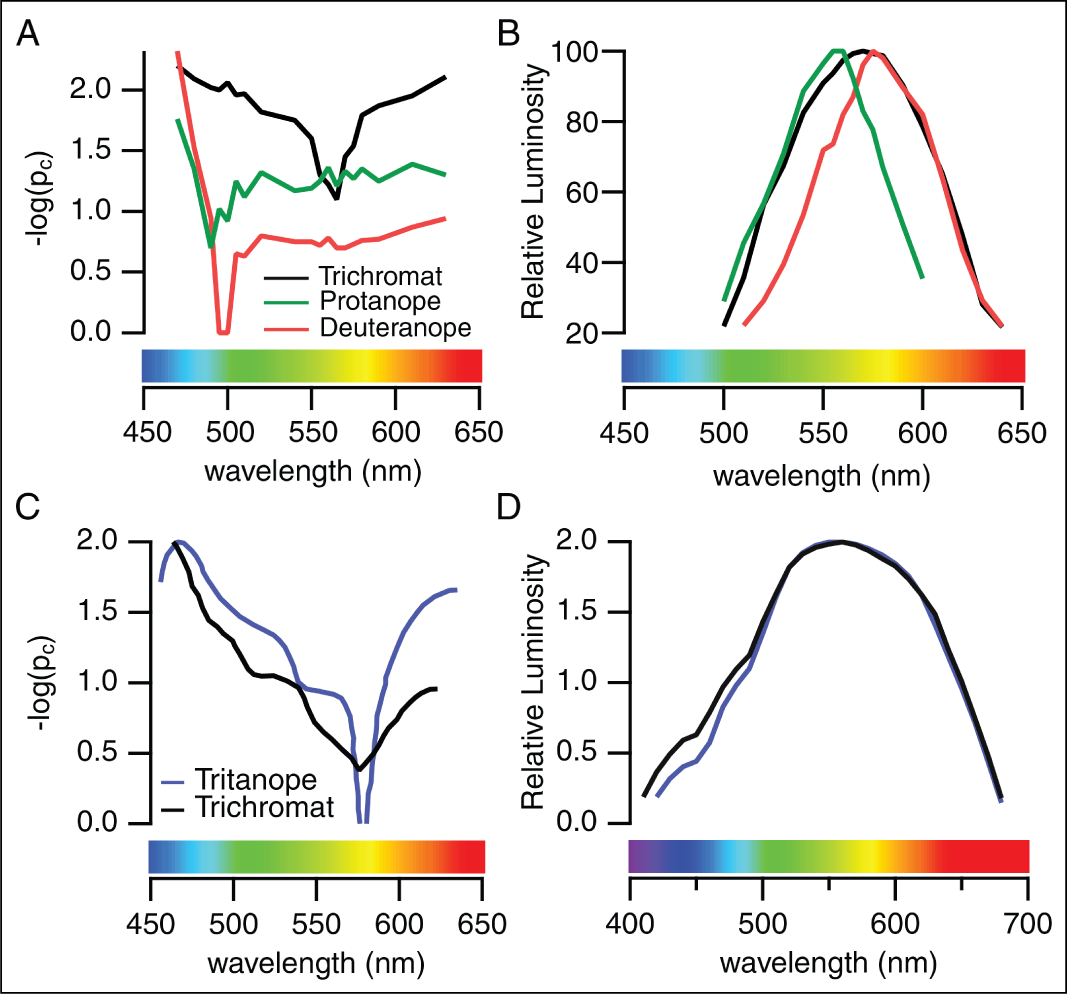
**(A)** Saturation discrimination and **(B)** spectral luminosity functions for trichromatic, deuteranopic and protanopic observers, from Chapanis (1944) [40]. **(C)** Saturation discrimination function for a tritanope and trichromat, from Cole (1966)[48]. **(D)** Average spectral luminosity functions for seven tritanopes compared to a trichromatic observer obtained under the same conditions, from Wright (1952) [49].

## 4. DISCUSSION

We find that spectrally-opponent midget ganglion cells can account for the saturation discrimination function measured using the “first step from white” paradigm. The midget ganglion cell models that successfully fit the “first step from white” paradigm could account for some, but not all, of the features of the scaling paradigm. Surprisingly, an S-OFF midget ganglion cell contribution was required to fully explain the “first step from white” saturation discrimination function. This is the first quantitative model to propose a role for the S-OFF midget ganglion cell in visual perception. Though the density of S-cones and S-OFF midget ganglion cells is far lower than L/M-cones and L/M-OFF midget ganglion cells, their contribution may be stronger because there is no opposing response from S-ON midget ganglion cells. An interesting avenue for future research will be to explore whether the S-OFF midget ganglion cells can explain any of the well-documented asymmetries in the S-cone ON and OFF pathways.

### A. Black and white percepts vs. the neural substrates for black and white

The saturation discrimination experiment does not directly involve sensations of black, however, the need for OFF midget ganglion cells to account for the “first step from white” paradigm suggests that the spectral sensitivity of white sensations is shaped by the neural substrates for both white and black. Indeed, some form of opponency is inherent in the anatomy and physiology of the ON and OFF pathways (reviewed by Westheimer, 2007 [39]) and opponency between black and white, is assumed in standard saturation models [4–6]. Importantly, our interpretation is meant to account for the saturation discrimination function and not for other measures of blackness and whiteness [37, 50–52].

### B. Interpreting the saturation discrimination experiment

The logic used to interpret the results of saturation discrimination experiments is as follows. The saturation of color depends not only on the strength of the hue component, but also the black-white component. If saturation is the relative portions of hue and black-white in a monochromatic light, then saturation experiments should measure how the relative amounts of hue and black-white vary as a function of wavelength. These variations as a function of wavelength reflect the spectral sensitivities of the neurons mediating hue and black-white sensations. It follows that saturation discrimination experiments can be interpreted as *a comparison between the spectral sensitivities of the neural substrates mediating our percepts of hue and black-white*.

Past models of saturation are based on the assumption that saturation represents the ratio of activity in the independent chromatic and achromatic channels, or the neural substrates of hue and black-white [4–6]. This can be expressed as an equation, like Hurvich & Jameson’s saturation coefficient:

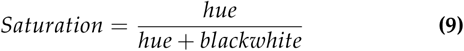

that proposes a relationship between the neural substrates for hue and black-white in defining color saturation. Regardless of the hue response to a given light, the perceived color saturation will also depend on the extent to which the neural substrates for black-white are simultaneously responding. For any given response from the neural substrates for hue, the color perceived will be less saturated if there is a large response from the neural substrates for black-white and more saturated if the black-white component of the total response is relatively small [34].

At each wavelength tested in the “first step from white” paradigm, the participant adjusted the intensity of the spectral light until a just-noticeable-difference (JND) in hue was perceived. Thus, at each wavelength tested, hue percepts are constant at 1 JND, a perceptual unit. Assuming our hue percepts correlate with activity in neural substrates for hue, we can propose that, for each wavelength tested in the “first step from white” paradigm, both the strength of the hue percept and the activity in the neural substrates for hue is effectively constant at 1 JND. This makes the results of the “first step from white” paradigm in Figure 5A a function of only one variable: black-white (Eq. (9)). The stronger the response from the neural substrates for black-white, the greater the intensity of the spectral light necessary to elicit a threshold hue response. The neural substrates for black-white respond to both the broadband light and the spectral light, and their spectral sensitivity determines the strength of their response to the spectral light. Thus, the “first step from white” paradigm can be interpreted as *revealing the underlying spectral sensitivity of the neural substrates for black-white*.

To put this interpretation in context, previous saturation models focused on the “first step from white” as a threshold change in saturation. For example, Hurvich & Jameson’s model was designed to find the hue and black-white (their chromatic and achromatic channels) responses necessary to give a just-noticeable-change in saturation. They assumed saturation was effectively constant at each wavelength tested and set Eq. (9) to be equal to 0.1, relative to 0 for a completely desaturated light [53]. As mentioned in the **Introduction**, when matched for brightness, each color has two components: hue and black-white and their relative strengths determine the perceived saturation. Here we propose that, in the case of the “first step from white” paradigms, the threshold change in saturation can be described more specifically as a threshold change in the hue component (i.e. a just-noticeable-difference in hue).

### C. Different saturation experiment paradigms measure distinct aspects of saturation

We now may be able to explain some of the variation between different measures of saturation discrimination, as shown in Figure 1. In one of the first studies of saturation discrimination, Purdy (1931) concluded the “first step from white” results are closely related to saturation, but do not completely match our perceptual experience of saturation [15]. Here, we reason they measure black-white instead.

The most accurate experiment for measuring our perceptual experience of saturation may be the results of saturation scaling experiments like Figure 1D because these tasks involved direct judgements of perceived saturation. The scaling experiments capture both the desaturation of yellows (~ 570 nm) and cyans (~ 490 nm) while Figure 1A emphasizes only the desaturation around 570 nm. We can reason that, unlike the “first step from white” experiments, scaling experiments engage both the neural substrates for hue and black-white. In other words, the results are a function of both variables in Eq. (9). Thus, a minimum in saturation can occur because the black-white response is relatively strong, or because the hue response is relatively weak. Comparison with opponent response functions [34, 35] indicate the latter is the case for the 490-500 nm region.

Our approach may also be able to explain the striking differences in saturation measured as the “first step from the spectrum” (Figure 1B) [15–17]. Like the “first step from white”, a threshold change in saturation is measured, however, in the “first step from the spectrum” paradigm, it is defined as a JND in white, not hue (see **Methods**). We can reason that this is a situation where black-white sensations are held constant and only hue sensations and the neural substrates of hue are predicted to vary. Thus, the “first step from the spectrum” paradigm measures a distinct process (the neural substrates of hue) and would not be expected to match the results in Figure 1A or Figure 1D.

Our model indicates that the spectral opponency provided by the midget ganglion cell’s center-surround receptive field is crucial for establishing the spectral sensitivity of downstream percepts. An interesting implication is that any experimental condition that minimizes contributions from spatial opponency will result in non-opponent spectral sensitivity. For example, spatial opponency weakens at high temporal frequencies [54, 55] and low light levels [56] so experiments involving these conditions are predicted to reveal spectral sensitivities like *V*(*λ*), the sum of L- and M-cones without evidence for spectral opponency.

### D. What is “achromatic”?

As discussed above, early saturation models, such as Hurvich & Jameson’s opponent process theory, represented saturation as the ratio of activity in chromatic and achromatic channels [4, 34]. Hurvich & Jameson’s achromatic channel refers to black, white and gray, or the absence of any hue sensations. However, vision scientists often associate “achromatic” with neurons that are spectrally non-opponent, summing the responses of different cone types, and unable to distinguish changes in wavelength from changes in intensity. Meanwhile, “chromatic” neurons are spectrally-opponent and compare the activities of different cone types.

Accordingly, Hurvich & Jameson (1955) assumed the mechanisms mediating black-white percepts (their “achromatic channel”) matched *V*(*λ*), the spectral luminosity function. Thus, “achromatic” has often been equated with luminance, *V*(*λ*), L+M-cone activity, and later, the activity of non-spectrally-opponent neurons, particularly the parasol ganglion cells [57]. We note in passing that there are other *L* + *M* ganglion cells in the primate retina with similar response properties to parasol ganglion cells [58] and *V*(*λ*) can also be obtained from the responses of *L − M* midget ganglion cells [59].

Saturation has proven difficult to account for by models assuming black and white percepts are mediated by neurons with spectral sensitivities matching *V*(*λ*). Figure 1A and Figure 1C show the saturation discrimination and luminosity functions for the same two observers [12, 13], which diverge in two key features. If the “first step from white” experiment measures the spectral sensitivity of the neural substrates for black-white, then models using *V*(*λ*) as a starting assumption may need to be revisited. The role of *V*(*λ*) is well-summarized by a review on luminance by Lennie and colleagues (1993): “V(*λ*) is a useful and well-established standard for photometry, but beyond that its significance for visual function is unclear” [59].

### E. Black-white sensations can be accounted for by spectrally-opponent midget ganglion cells

Increasing evidence supports the idea that midget ganglion cells mediate high resolution achromatic spatial vision (reviewed by Lennie & Movshon (2005) [60]). As discussed above, their role in spatial vision underlies their ability to encode black and white percepts, because these are defined at the borders of objects [41, 42, 61]. The center-surround receptive fields of ON and OFF midget ganglion cells are perfectly suited for this task [43] - they encode increments and decrements in spatial contrast, respectively, by encoding the photon catch in the center cone relative to the photon catch in surrounding cones.

However, their role in encoding black and white percepts has been difficult to reconcile with their classification as “chromatic” neurons (reviewed in Patterson et al., 2019 [9]). From an individual midget ganglion cell’s spike output, it is impossible for a downstream neuron to distinguish whether the ganglion cell responded to an edge defined by a change in intensity or spectral contrast. Our intuition is that the midget ganglion cell’s spectral and spatial opponency must be separated downstream, so that responses to changes in wavelength result in hue percepts while responses to changes in intensity result in black and white percepts. Models proposing midget ganglion cells mediate black-white typically assume this downstream separation (or “demultiplexing”) must occur [8]. However, the results of saturation discrimination experiments are contrary to our intuition - if changes in wavelength only resulted in hue percepts and changes in intensity only resulted in black-white percepts, saturation would not vary as a function of wavelength and the results of the saturation scaling experiment in Figure 1D would be a constant 100% saturation (pure hue with 0% white percepts).

Here we demonstrated the saturation discrimination function can be accounted for by linear combinations of the outputs of midget ganglion cells, without assuming downstream “demultiplexing” occurs. In light of this result, we might consider an alternative: that the five types of ON and OFF midget ganglion cells illustrated in Figure 3 are “labeled lines” for white and black, or more accurately, edges defined by increments and decrements in spatial contrast, respectively, and that other ganglion cells mediate hue perception [10, 62, 63].

An inability to distinguish between edges defined by intensity and spectral contrast is not a problem if the midget ganglion cell’s only function is only to encode edges (and not hue). After all, an optimal edge detector should be able to detect all edges, including the equiluminant ones commonly found in natural scenes [64]. A consequence is that the difference in spectral tuning between center and surround receptive fields will endow the edge detection that defines our percepts of black and white with a clear spectral sensitivity. For example, an ON midget ganglion cell will respond more to an edge between a broadband light and a 570 nm light than an edge between a broadband light and a 450 nm light (Figure 4D, 5B, 5D), thus signaling greater spatial contrast and a stronger white sensation for the 570 nm edge. Put simply, the striking wavelength-dependence of saturation can be explained at the level of the retinal output by spectrally-opponent edge detectors.

## 5. CONCLUSIONS

In a classic study on saturation discrimination, Purdy (1931) remarked, “No theory of color vision, however, can be regarded as adequate if it does not account for the laws of saturation as well as those of hue.”[15]. We demonstrate that spectrally-opponent midget ganglion cells provide the most accurate account to date of the physiological mechanisms underlying saturation.

## 6. ACKNOWLEDGMENTS

Portions of this work were presented at the International Color Vision Society meeting in 2019.

